# Estimation of *in vivo* constitutive parameters of the aortic wall: a machine learning approach

**DOI:** 10.1101/366963

**Authors:** Minliang Liu, Liang Liang, Wei Sun

**Author notes:** For correspondence: Wei Sun, Ph.D., The Wallace H. Coulter Department of Biomedical Engineering, Georgia Institute of Technology and Emory University, Technology Enterprise Park, Room 206, 387 Technology Circle, Atlanta, GA 30313-2412.

## Abstract

The patient-specific biomechanical analysis of the aorta demands the *in vivo* mechanical properties of individual patients. Current inverse approaches have shown the feasibility of estimating the nonlinear, anisotropic material parameters from *in vivo* image data using certain optimization schemes. However, since such inverse methods are dependent on iterative nonlinear optimization, these methods are highly computation-intensive, which may take weeks to complete for only a single patient, inhibiting rapid feedback for clinical use. Recently, machine learning (ML) techniques have led to revolutionary breakthroughs in many applications. A potential paradigm-changing solution to the bottleneck associated with patient-specific computational modeling is to incorporate ML algorithms to expedite the procedure of *in vivo* material parameter identification. In this paper, we developed a ML-based approach to identify the material parameters from three-dimensional aorta geometries obtained at two different blood pressure levels, namely systolic and diastolic geometries. The nonlinear relationship between the two loaded shapes and the constitutive parameters are established by a ML-model, which was trained and tested using finite element (FE) simulation datasets. Cross-validation was used to adjust the ML-model structure on a training/validation dataset. The accuracy of the ML-model was examined using a testing dataset.

## 1. INTRODUCTION

With advances in medical imaging modalities and computation power, numerical simulations of the cardiovascular structure such as the aorta, which utilizes the patient-specific three-dimensional (3D) geometry, have been increasingly popular (Taylor and Figueroa, 2009). Yet, the difficulty in obtaining patient-specific elastic properties of the aortic wall from *in vivo* images has been one of the biggest obstacles in front of patient-specific biomechanical analysis. This has motivated recent efforts to develop inverse methods for estimating the *in vivo* material properties of the aortic wall on a patient-specific basis. In these methods, deformations and boundary conditions are used to inversely estimate the material parameters of a particular constitutive model. However, the complex 3D shapes and nonlinear and anisotropic constitutive behavior make this task challenging.

To avoid computational complexity, some studies suggested the use of simplifications in material models and/or geometries. For example, (Liu and Shi, 2009), (Zhang et al., 2017) and (Franquet et al., 2013) identified linear elastic material parameters. By assuming a perfect cylindrical shape of the arteries, (Schulze-Bauer and Holzapfel, 2003) identified Fung-type material parameters, (Masson et al., 2011; Masson et al., 2008; Olsson and Klarbring, 2006; Stålhand, 2009) estimated material parameters using the constitutive model proposed in (Holzapfel et al., 2000), and (Smoljkić et al., 2015) estimated the Gasser–Ogden-Holzapfel (GOH) model (Gasser et al., 2006) parameters. (Liu et al., 2012) also determined the modified Moony-Rivlin parameters of the carotid artery from 2D slices reconstructed from MRI.

To fully exploit the 3D geometries reconstructed from medical image data, the current *in vivo* material parameter estimation methods largely rely on optimization schemes. In these optimization-based inverse methods, an objective/error function is built upon the difference between predicted and image-measured physical fields (e.g. stress/strain/displacement), and then the constitutive parameters are iteratively adjusted to ensure the objective function is minimized. Specifically, with an initial guess of the constitutive parameters, (1) the unloaded configuration is recovered, (2) the physical field in the loaded state is computed by applying *in vivo* loading and boundary conditions, and (3) the constitutive parameters are iteratively fine-tuned by a nonlinear optimization algorithm until the predicted physical field matches with the image-measured physical field. This optimization process yields the optimal constitutive parameters. Using finite element (FE) updating schemes, (Wittek et al., 2016; Wittek et al., 2013) developed two methods to determine GOH material parameters of the human abdominal aorta from *in vivo* 4D ultrasound data. However, numerous iterations were needed to reach the optimal solution, resulting in long computing time of 1~2 weeks. Such high computational cost could inhibit the practical use of these methods, particularly in a clinical setting requiring rapid feedback to clinicians. To this end, our group has recently proposed two optimization-based methods to expedite the estimation process. The multi-resolution direct search (MRDS) approach (Liu et al., 2018) was designed to improve the searching algorithm, and the computation time was reduced to 1~2 days with similar CPU and memory. In the stress-based inverse approach (Liu et al., 2017), the computationally-expensive FE simulations were avoided by building the objective function upon stresses, and the optimization was completed in approximately 2 hours. However, the optimization-based methods are inherently limited by their iterative nature, and any further improvement of speed becomes difficult.

Recently, machine learning (ML) techniques, particularly deep learning (DL) (LeCun et al., 2015; Litjens et al., 2017; Shen et al., 2017), have garnered enormous attention in the field of artificial intelligence, leading to revolutionary breakthroughs in many applications (Hannun et al., 2014; He et al., 2015, 2016; Kokkinos, 2016; Krizhevsky et al., 2012; LeCun et al., 2015; Taigman et al., 2014; Wu et al., 2016). ML-models are capable of establishing complex and nonlinear relationship between inputs and outputs. A potential paradigm-changing solution to the bottleneck associated with patient-specific computational modeling is to incorporate ML algorithms to expedite the procedure of *in vivo* material parameter identification. By designing and training a ML-model on a large dataset, it can automatically produce the required outputs (constitutive parameters) directly from necessary inputs (multi-phase aorta shapes), without the need for costly iterative schemes. Once trained, the ML-model can instantaneously predict the material parameters.

**Figure 1.**
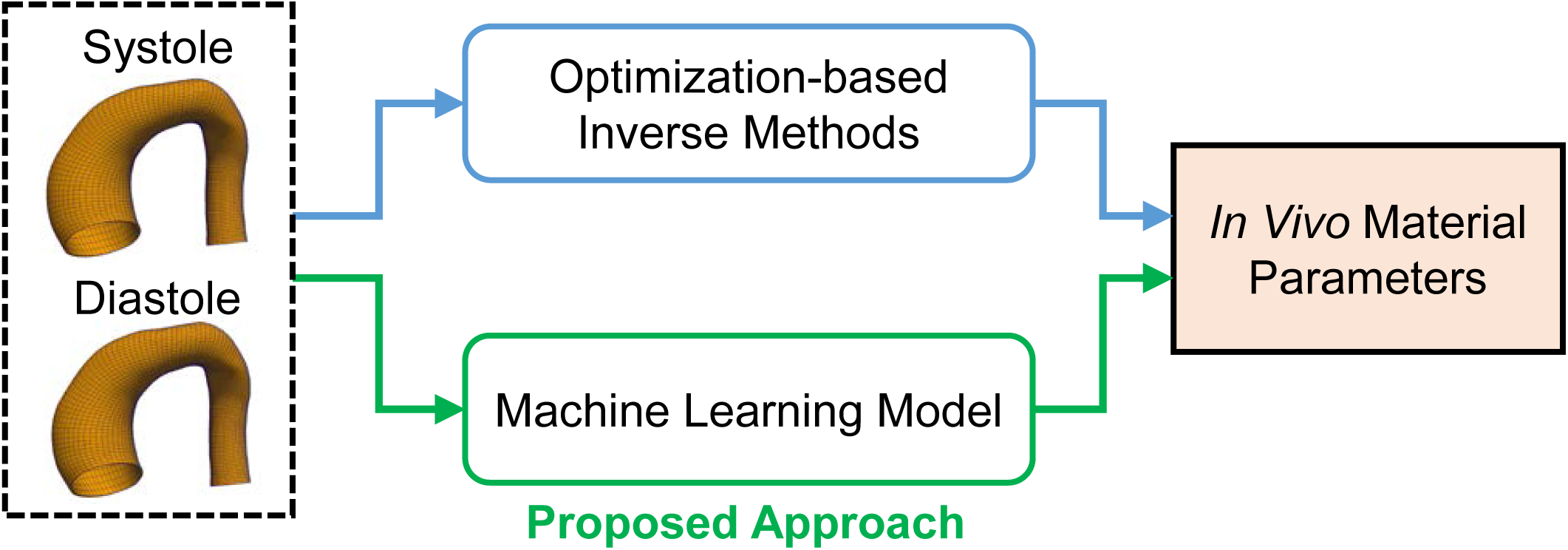
The proposed machine learning (ML) approach.

In this paper, we developed a ML-based approach to identify the material parameters of the GOH constitutive model. As depicted in Figure 1, the inputs to this ML-model are aorta geometries at two distinct blood pressure levels, namely systolic and diastolic geometries, which were also used by our previous optimization-based inverse approaches. The nonlinear relationship between the two loaded shapes and the constitutive parameters are established by a ML-model, consisting of an unsupervised shape encoding module (principal component analysis) and a supervised nonlinear mapping module (neural network). The datasets for training, validation and testing were generated from FE simulations. Cross-validation was used to adjust the neural network structure. The accuracy of the ML-model was examined using a testing dataset.

## 2. METHODS

### 2.1 Constitutive model

A strain invariant-based fiber reinforced hyperelastic formulation based on the work of (Gasser et al., 2006) was used to model the constitutive relations of the aortic wall. The deformation gradient **F** can be multiplicatively decomposed into

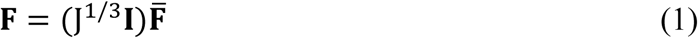

where J is the determinant of **F**, and **I** is the identity tensor. 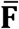 represents the volume-preserving (isochoric) part of the deformation gradient, while (J^1/3^**I**) represents the volumetric part. The right Cauchy-Green tensor ***C*** and its isochoric counterpart 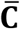 is defined as

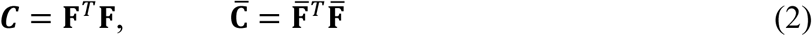

The isochoric strain invariants 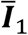 and 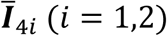 are defined using

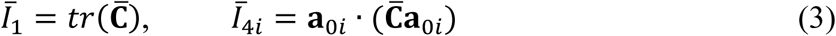

where vectors **a**_01_=(cos*θ*, sin*θ*, 0) and **a**_02_=(cos*θ*, −sin*θ*, 0) characterize the two mean fiber directions in the reference configuration, they are defined in the circumferential direction. Thus, 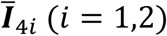 are equal to squares of the stretches in these directions. The total strain energy function Ψ can be additively split into isochoric isotropic 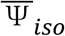, isochoric anisotropic 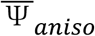 and volumetric Ψ*_vol_* parts, according to

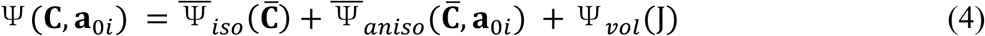

The isotropic matrix material is characterized by the Neo-Hookean strain energy

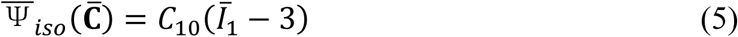

where *C*_10_ is a material parameter to describe the matrix material. The isochoric anisotropic term is given by

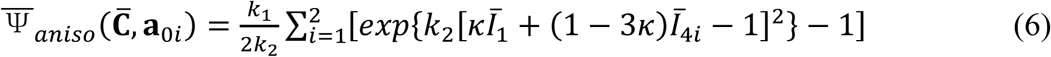

Where *k*_1_ is a positive material parameter that has the same unit of stress, while *k*_2_ is a unitless material parameter. *κ* ∈ (0,1/3) describes the dispersion of fibers. Finally, the volumetric term is defined by

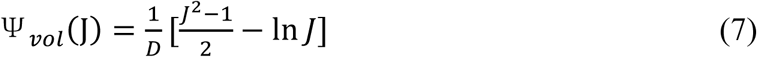

Where *D* is a constant that enforces material incompressibility and it is fixed to 5 × 10^−4^. Thus, the aortic wall tissue is characterized by five constitute parameters {*C*_10_,*k*_1_,*k*_2_,*κ*,*θ*}. The task for the ML-model is to identify the five constitute parameters given the two-phase aorta shapes.

### 2.2 Generating the training/validation dataset and the testing dataset

The proposed ML model will establish a mapping between the inputs (geometries) to the outputs (material parameters) based on example input-output pairs. Each input-output pair consists of two geometries and the corresponding material parameters. To fine-tune the ML-model for optimal performance, cross-validation is used in the training phase, where the input-output pairs are partitioned into two subsets, called training set and validation set. The ML-model is trained on the training set, and its performance is assessed using the validation set. After the training phase, the accuracy of the ML-model is evaluated on a new set of input-output pairs, i.e., the testing set.

In this approach, the datasets are gathered from FE simulations. Using statistical modeling methods, representative material parameters are generated from 65 sets of experimentally-derived material parameters, and representative virtual aorta geometries at one physiological phase (systole) are generated from 3D CT images of 25 real patients. The diastolic aorta geometries are determined from FE simulations using the virtual systolic geometries and the representative material parameters. Finally, the training/validation dataset and the testing dataset consist of 15366 and 225 input-output pairs, respectively. The detailed procedures to generate the datasets are presented in the following subsections.

#### 2.2.1 Sampling the material parameter space

In previous studies (Martin et al., 2013; Pham et al., 2013), seven-protocol biaxial tension experiments were carried out on a total of 65 aneurysmal patients, and five material parameters of the GOH model were determined by fitting the experimentally-obtained stress-strain curves. The material of a patient was represented by a vector, ***y***^(^*^i^*^)^ (*i* = 1,2,…,65), with its five components corresponding to five GOH parameters, and the set ***Y*** contained all the 65 vectors. These vectors are visualized in the material parameter space in Figure 2, which shows that these experimentally-derived parameters are highly clustered in certain regions. To uniformly sample the material parameter space, a convex hull of the experimentally-derived parameters was built. The convex hull is essentially a set comprised of convex combinations of all vectors of ***Y***,

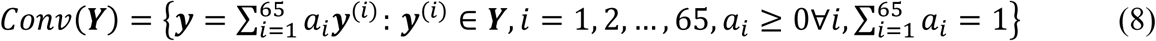

where ***y*** is a vector in the convex hull, and *a_i_* (*i* = 1,2,…,65) are nonnegative coefficients that sum up to 1. Next, samples were draw from a uniform distribution inside the convex hull using the Gibbs sampler (Geman and Geman, 1984). 125 and 15 samples were generated for the training/validation set and the testing set, respectively, as shown in Figure 2.

**Figure 2.**
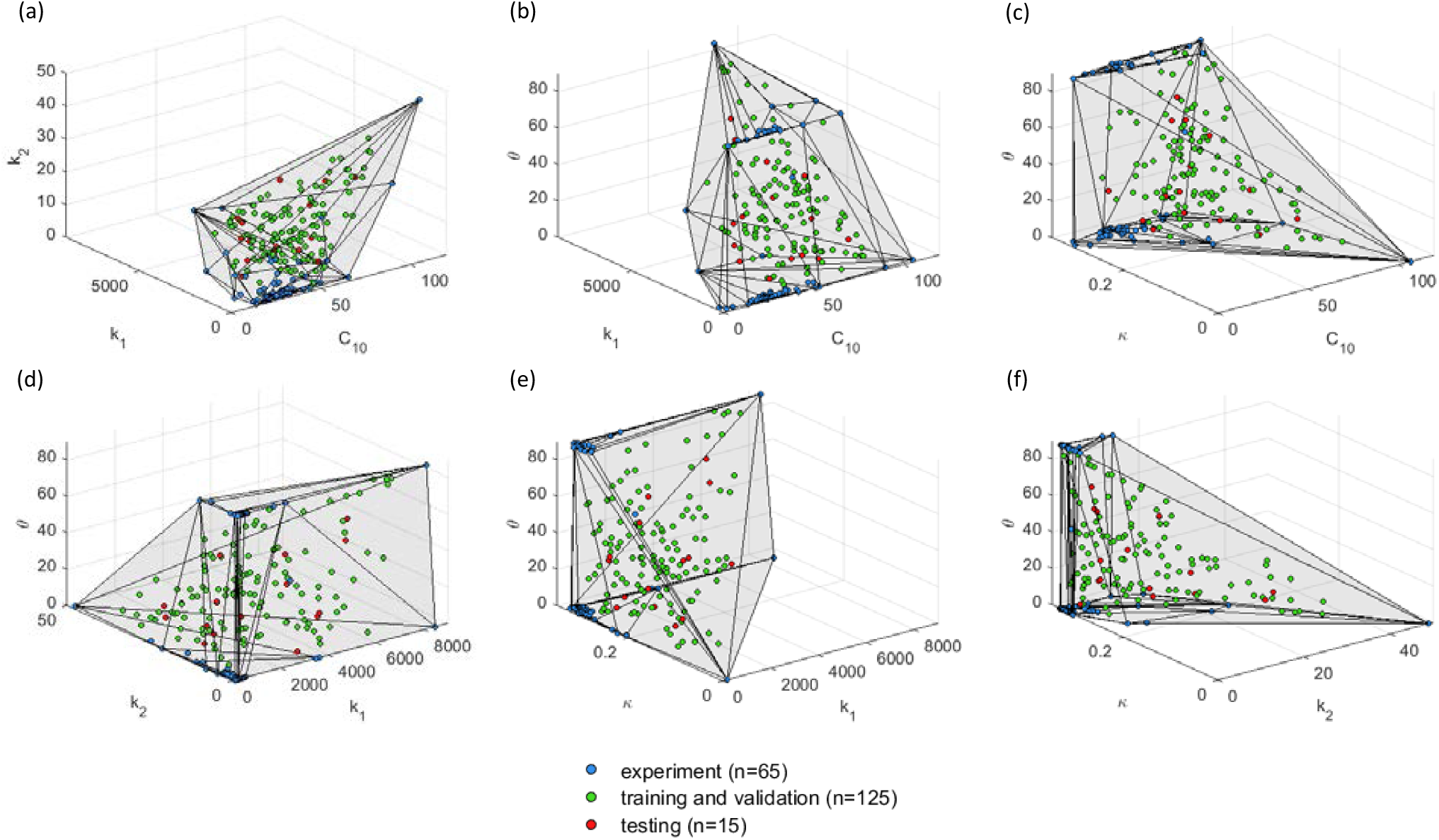
Datasets projected in 3D material parameter subspaces. The convex hull is plotted in the 3D subspaces for illustrative purpose.

#### 2.2.2 Obtaining the virtual aorta geometries at the systolic phase

From a previous study (Martin et al., 2015), 3D CT images of 25 real patients who underwent elective repair were collected from Yale-New Haven Hospital. A statistical shape model (SSM) was built from the 25 real aorta shapes at the systolic phase in a subsequent study (Liang et al., 2017). The systolic geometries can be represented by a set of SSM parameters {*C*_1_,*C*_2_,*C*_3_}. Please refer to our previous paper (Liang et al., 2017) for further details.

For the training and validation dataset, a total number of 125 virtual aorta shapes at systolic phase were obtained by sampling the SSM parameter space with equally spaced points in the range of −2 to 2, i.e., within 2 standard deviations of the mean shape, as shown in Figure 3. Similarly, for the testing dataset, the SSM parameter space was sampled within 1.5 standard deviations of the mean shape. Hence, 15 systolic shapes were obtained for the testing dataset. The resulting samples are plotted in the SSM parameter space in Figure 3.

**Figure 3.**
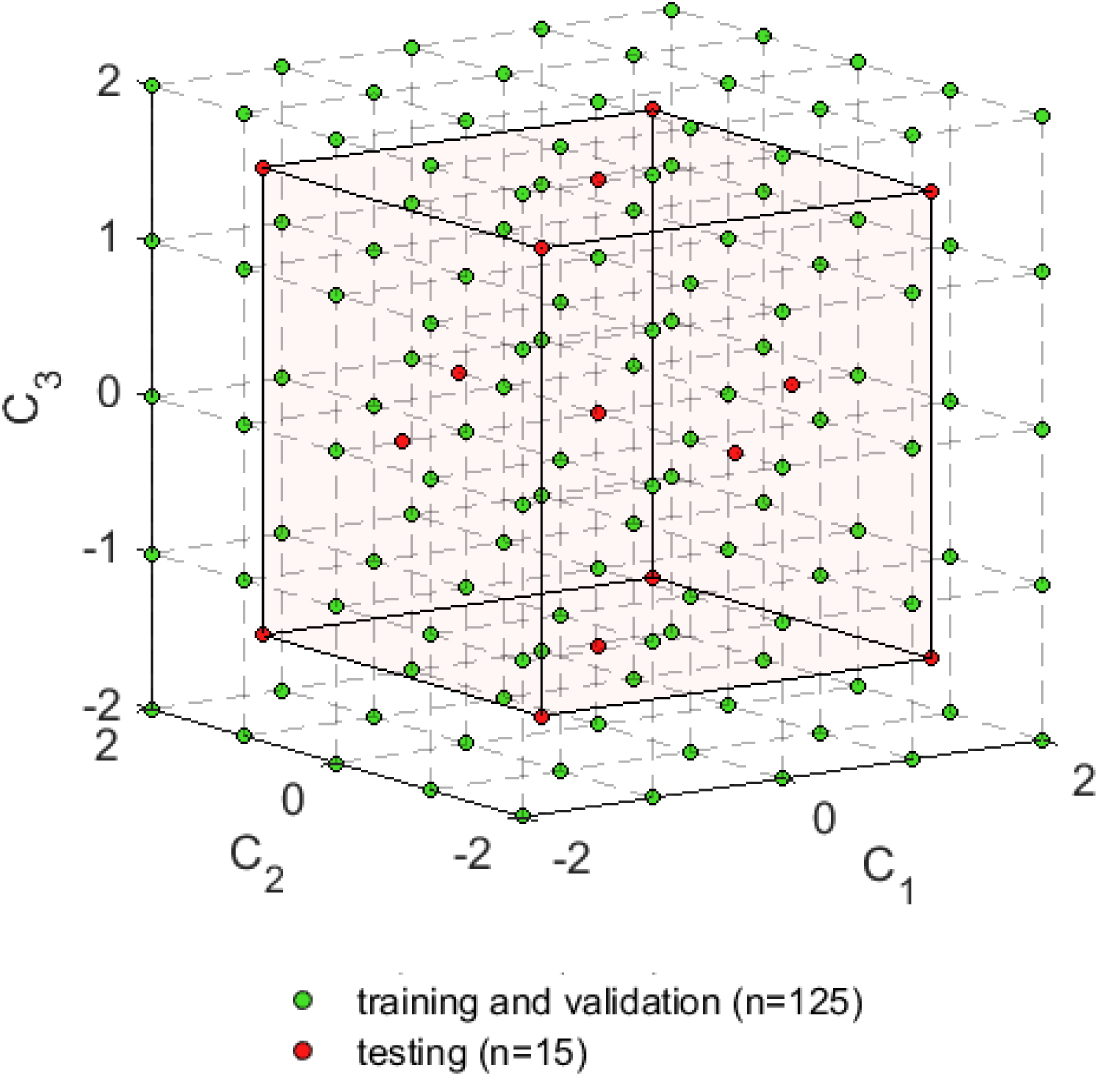
Sampling the SSM parameter spaces.

#### 2.2.3 Generating the virtual aorta geometries at the diastolic phase using FE simulations

The virtual aorta geometries obtained from the SSM parameter space were at the systolic phase, which should be in equilibrium with the systolic physiological load. Therefore, the generalized pre-stressing algorithm (GPA) (Weisbecker et al., 2014) was utilized to incorporate the pre-stress induced by the systolic pressure (16 kPa). In the GPA algorithm, the total deformation gradient ***F****_t_* is stored as a history variable for each integration point. The ***F****_t_* is updated based on the incremental deformation gradient Δ***F*** resulting from the prescribed loading and boundary conditions.

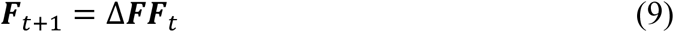

The incremental deformation gradient Δ***F*** resulting from the systolic pressure is iteratively applied to the virtual systolic geometries and stored in ***F****_t_*. However, as illustrated in the original paper (Weisbecker et al., 2014), the resulted equilibrium configuration slightly deviate from the original configuration, depending on the step size. The systolic geometries at the equilibrium configurations were used by the ML-model in the subsequent sections. Next, using a set of material parameters, the virtual aorta geometries at the diastolic phase were determined by depressurizing the systolic geometries to the diastolic phase (10 kPa).

For the training and validation set, given one of the 125 shapes at the systolic phase and one of the 125 sets of material parameters, the virtual aorta geometry at the diastolic phase was determined through FE simulation with GPA. As shown in Figure 4, if the FE simulation converges, the input-output pair (systolic and diastolic geometries and a set of material parameters) is collected for training/validation. As a result, 15366 sets of geometries with known material parameters were obtained. Similarly for the testing set, 225 input-output pairs were generated from 15 systolic geometries and 15 sets of material parameters.

The GPA algorithm was implemented in ABAQUS user subroutine UMAT. In the FE simulations, C3D8H solid elements were used, and pressure was applied uniformly to the inner surface of the aorta models. The boundary nodes of the aorta models, i.e. the proximal and distal ends of the model, were only allowed to move in the radial direction defined by the local coordinate system.

**Figure 4.**
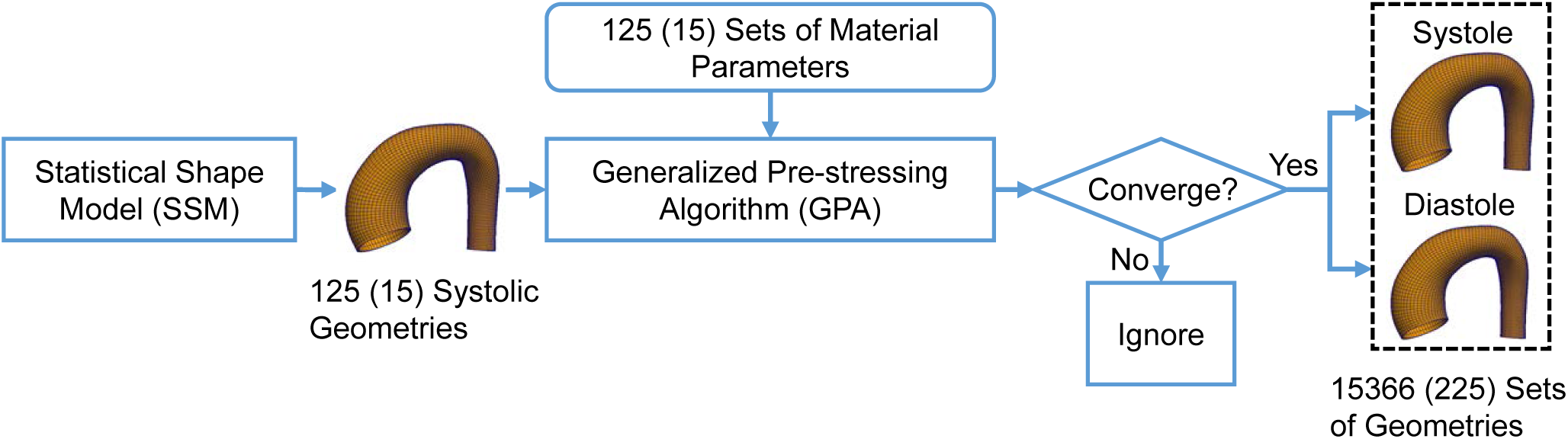
The procedure to generate aorta geometries at systole and diastole. The number in the parenthesis indicates the testing dataset.

### 2.3 The machine learning model

The machine learning model consists of an unsupervised shape encoding module and a nonlinear mapping module. The systolic and diastolic shapes are encoded by shape codes. The nonlinear mapping between the shape codes and the material parameters is established by a neural network.

#### 2.3.1 Shape encoding

3D geometries are usually represented by polygon meshes. A shape corresponds to a long vector ***X*** of nodal coordinates. However, directly linking the shape ***X*** to the material parameters by a neural network, although possible, can lead to a large number of undetermined parameters which needs a very large training dataset. A compact representation (i.e. shape code) of a shape can be obtained in a shape encoding procedure. The principal component analysis (PCA) (Webb and Copsey, 2011) is widely adopted as a shape encoding method and an unsupervised learning technique for dimensionality reduction, in which the original data can be well approximated by a linear combination of a few principal components. Thus, a shape *X* can be represented by

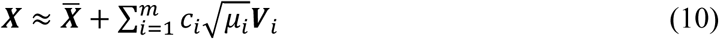

where *X* is the mean shape, ***V****_i_* and *μ_i_* are the eigenvectors (i.e. modes of shape variations) and eigenvalues of the covariance matrix, respectively. *m* is the number of modes used for approximation. The shape code {*c_i_*,*i* = 1,…,*m*} can be obtained by

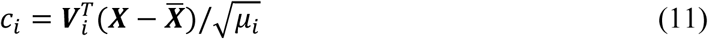

where 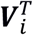 is the transpose of the column vector ***V****_i_*. The first 12 modes (*m*=12) were retained for both the systolic and diastolic shape encoding, with the average PCA approximation error being less than 0.1%. Note that as mentioned in Section 2.2.3, the systolic geometries from GPA were slightly different from the original configuration from the SSM, and therefore 3 modes are not enough to capture the shape variations for systolic geometries. We denote the systolic shape code as *α_i_*, diastolic shape code as *β_i_*, (*i* = 1,2,…,12).

#### 2.3.3 Nonlinear mapping

The nonlinear mapping module will map the shape codes of the two input shapes to the five material parameters, which is equivalent to establishing five nonlinear functions

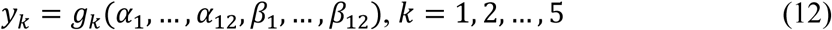

The inputs are the shape codes for diastolic and systolic geometries *α_i_* and *β_i_*, (*i* = 1,2,…,12). The outputs are *y_k_* (*k* = 1,2,…,5), correspond to the five material parameters {*C*_10_,*k*_1_,*k*_2_,*κ*,*θ*}.

As shown in Figure 5, a neural network is constructed as the nonlinear mapping module. It consists of feedforward fully-connected units (neurons). Each unit has multiple inputs and a single output. For the *j*th unit of the *i*^th^ layer, a linear combination of the input vector ***z****^i^* (with weight vector 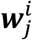 and offset 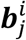) is computed as

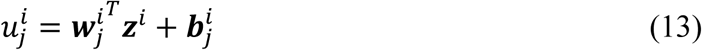

where the superscript *i* denote the index of layer, and subscript *j* denote the index of unit in the layer. ***z****^i^* is a column vector of 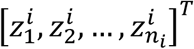, and *n_i_* is the number of units in the *i*^th^ layer. The weighted sum 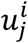 is nonlinearly transformed into the output 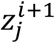 using an activation function.

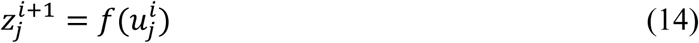

The softplus (Dugas et al., 2000) activation function was used, which is given by

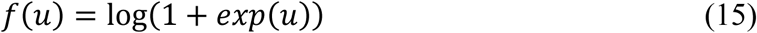

This function is a smooth version of the rectified linear unit (ReLU) (Glorot et al., 2011; Hahnloser et al., 2000). As demonstrated in the discussion section, other activation functions can lead to large testing errors in our application. The neural network has two hidden layers with the same number of softplus units, and the output layer has 5 softplus units.

The neural network was implemented using Tensorflow (Abadi et al., 2015). The inputs and outputs were normalized using the maximum absolute value of each dimension. Consequently, the normalized shape codes are within the range of −1 to 1, and the normalized material parameters are within the range of 0 to 1. The mean squared error (MSE) was used as the loss function

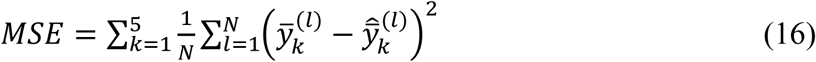

where *l* is the index for an input-output pair, *N* is the total number of input-output pairs, 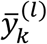 and 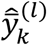 represent the *k*^th^ actual and predicted normalized material parameter, respectively. After the nonlinear mapping, the predicted material parameters were rescaled to its original range. The parameters of the neural network were obtained through the Adamax optimization algorithm (Kingma and Ba, 2015). For detailed theories, please refer to (Goodfellow et al., 2016).

**Figure 5.**
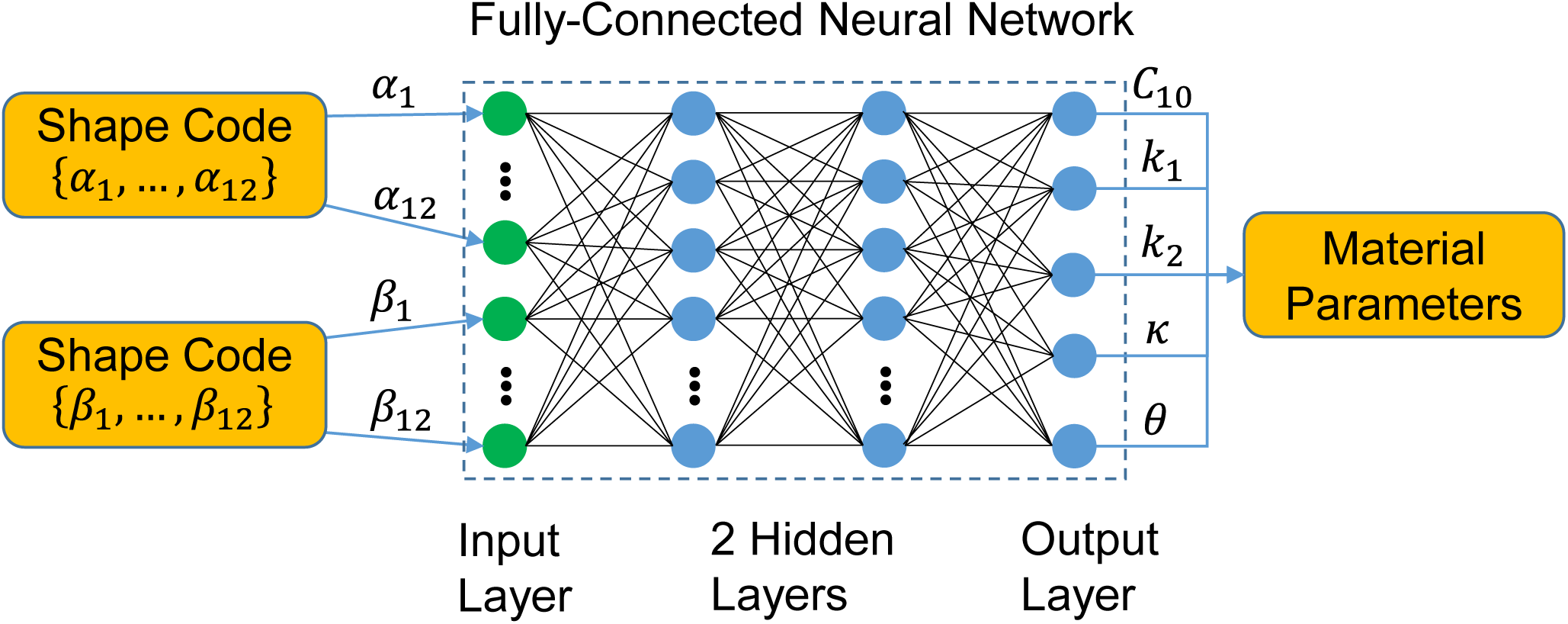
The neural network for mapping the shape codes to the material parameters. The green dots represents the input layer, and the blue dots represent the softplus units in the hidden layers and the output layer of the neural network.

### 2.4 Training, adjusting and testing the ML-model

The unsupervised shape encoding module (PCA) was built only upon the training and validation sets. For shapes in the testing set, the shape codes were obtained from Eqn. (4) using 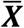, ***V****_i_* and *μ_i_* computed from the training and validation set.

Using the training/validation dataset, the performance of the nonlinear mapping module was assessed through leave-one-out (LOO) cross-validation, and the neural network structure was fine-tuned. As depicted in Figure 6, in each round of the LOO cross-validation, the data was split into a training set and a validation set, according to the material parameters. We pick one set of material parameters (and its corresponding geometries) from the 125 sets from Section 2.2.1 as the validation set, and train the neural network on the remaining 124 sets (and its corresponding geometries). An averaged error was obtained after repeating this procedure for all of the 125 sets of material parameters. Hence, the training set never contains the material information used in the validation. Next, the number of units in each layer was adjusted to minimize the average error in the LOO cross-validation. The final network contains 256 units for each of the two hidden layers.

**Figure 6.**
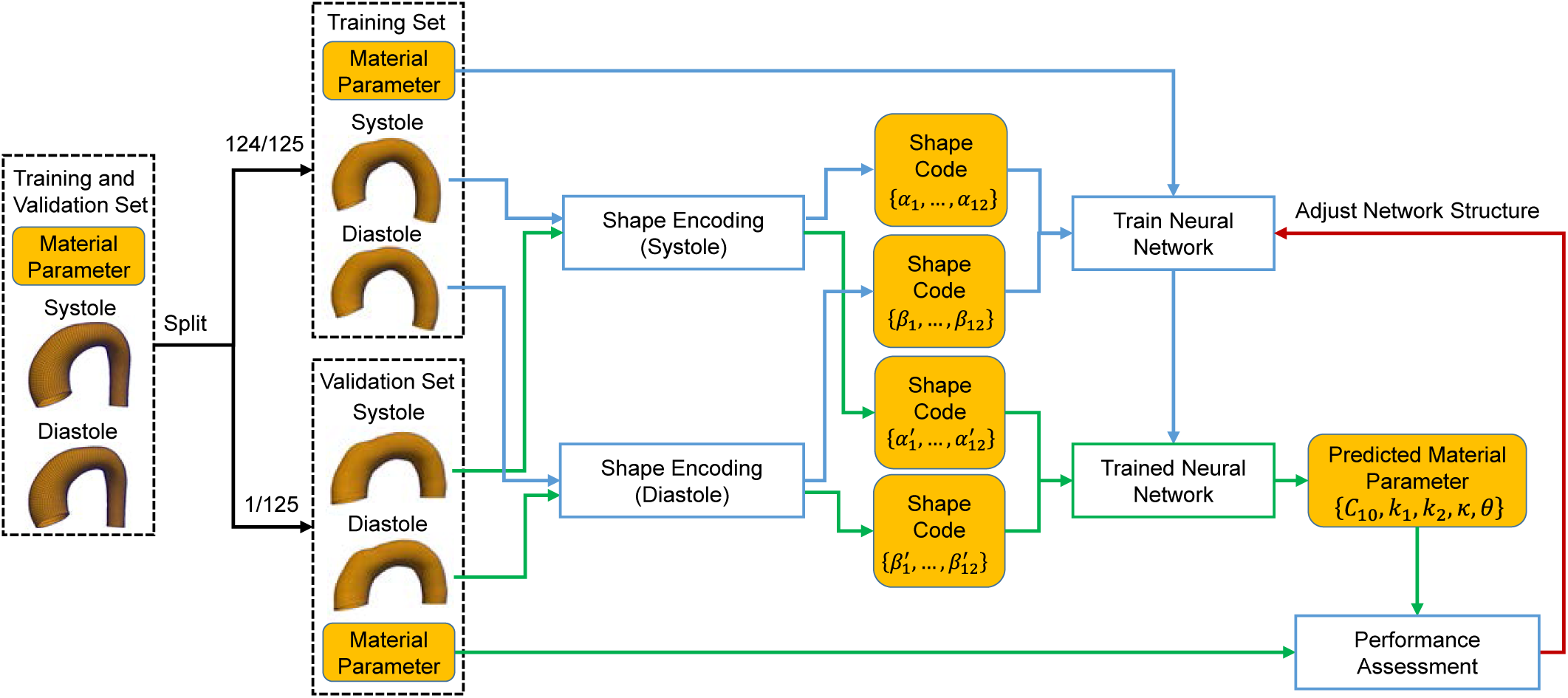
Adjusting the network structure using the leave-one-out (LOO) cross-validation.

To evaluate the ML-model, that is, to examine how accurate the prediction is compared to FE simulation data, the ML-model was trained on the training/validations set and then the material parameters were predicted using shapes in the testing set. The predicted material parameters were compared to the actual parameters in the testing set.

**Figure 7.**
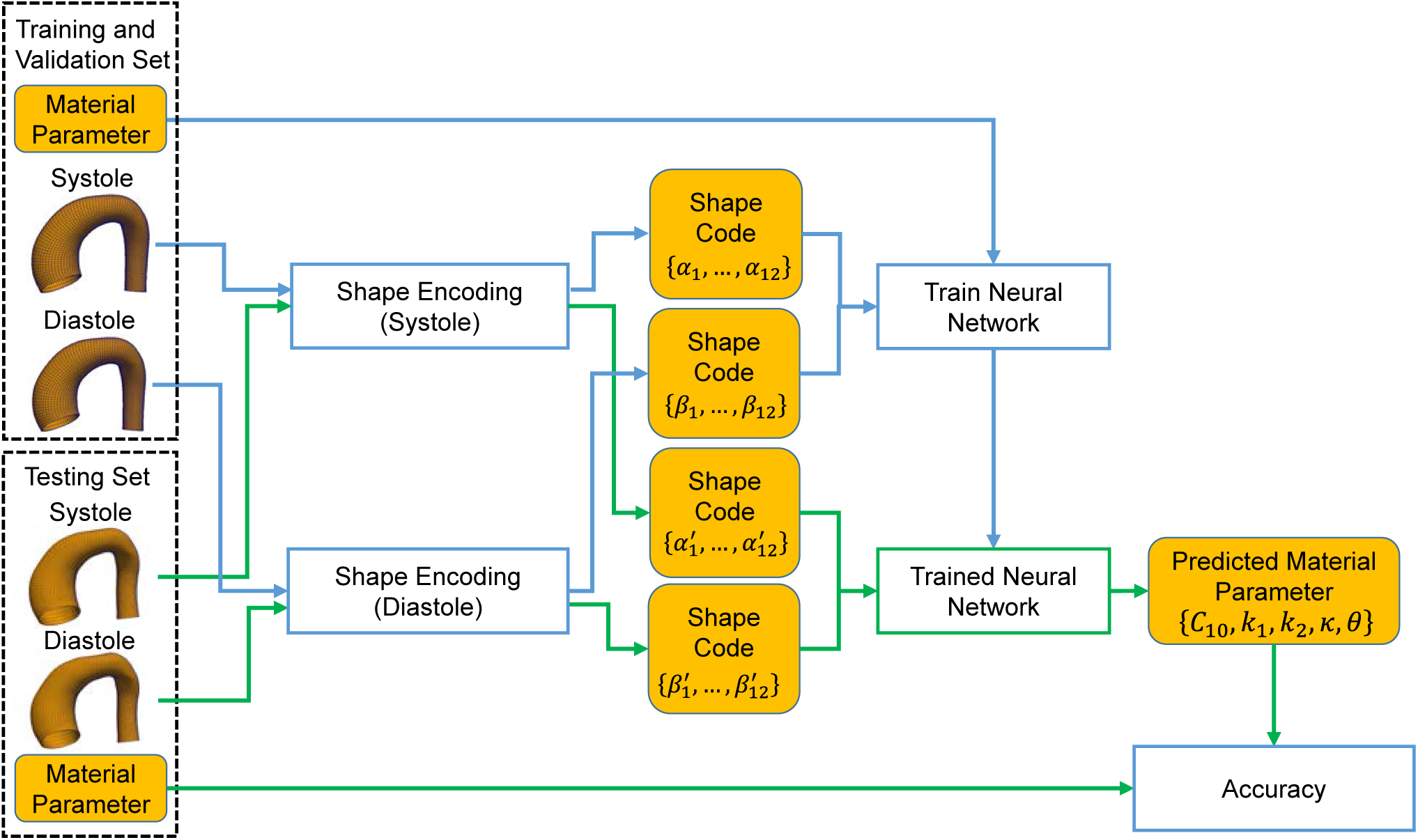
Evaluating the accuracy using the testing dataset.

## 3. RESULTS

Given a pair of geometries as the inputs, the trained ML-model can output the material parameter within one second on a PC with 3.6GHz quad core CPU and 32GB RAM. The actual versus predicted material parameters are shown in Figure 8. The discrepancy between the actual and predicted material parameters in the testing set was quantified by normalized mean absolute error (NMAE) and the normalized standard deviation of absolute error (NSTAE). The absolute error (AE) for the *k*^th^ material parameter is defined by

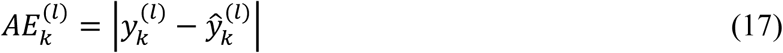

where index *l* and *k* are the same as Eqn.(17), 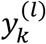 and 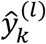 represent the *k*^th^ actual and predicted material parameter, respectively. The NMAE of the *k*^th^ material parameter is defined by

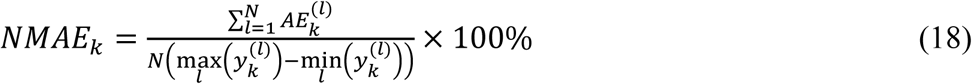

where *N* is defined in Eqn.(16). And the NSTAE of the *k*^th^ material parameter is

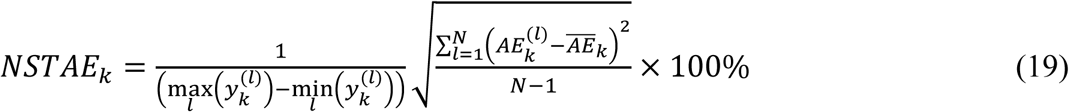

where 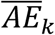 is the averaged absolute error for the *k*^th^ material parameter. The NMAE and NSTAE for each material parameter in the testing set are reported in Table 1. The errors indicate that the ML-predicted material parameters are in good agreement with the actual material parameters.

**Table 1.**
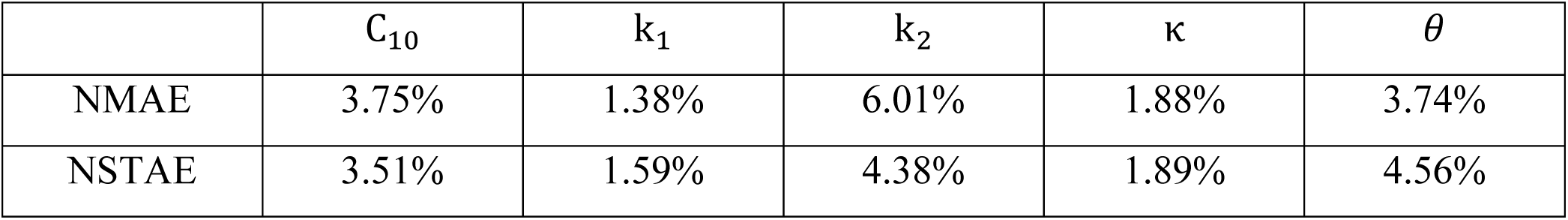
NMAE and NSTAE of the five material parameters in testing set.

**Figure 8.**
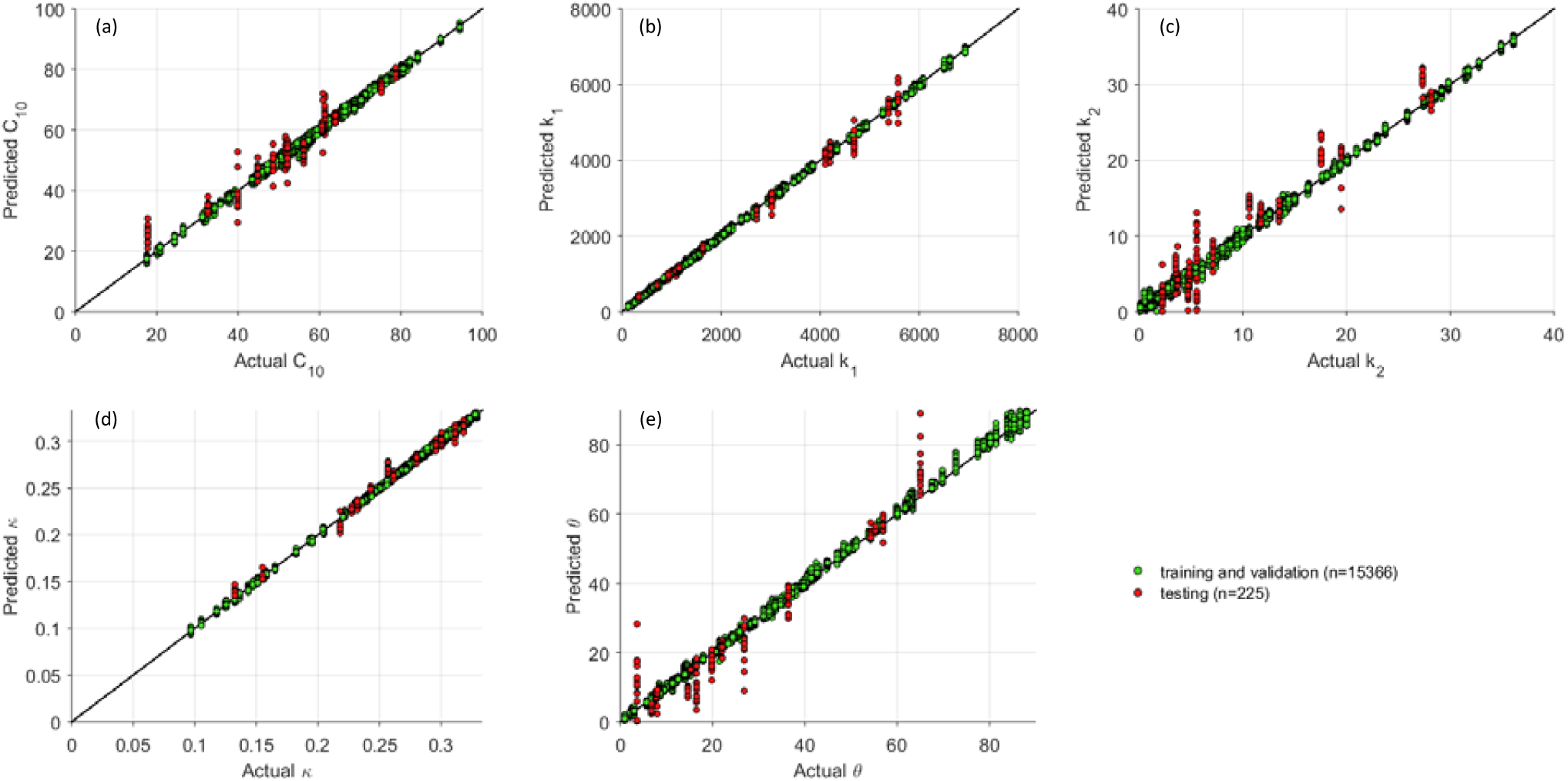
The actual and predicted material parameters.

To further evaluate the estimation results, stress-stretch curves were plotted by simulating stretch-controlled biaxial tension in MATLAB by assuming the tissue is loaded in the plane stress state and the material is incompressible. We use *σ*_1_ and *λ*_1_ to denote the circumferential stress and stretch, *σ*_2_ and *λ*_2_ to denote the longitudinal stress and stretch. The simulations were based on the following 3 protocols: (1) in the circumferential strip biaxial tension, fixing *λ*_2_ = 1 while increasing *λ*_1_; (2) in the equi-biaxial tension, keeping the ratio *λ*_1_/*λ*_2_ = 1; (3) in the longitudinal strip biaxial tension, fixing *λ*_1_ = 1 while increasing *λ*_2_. In total, six stress-stretch curves are generated for each set of constitutive parameters.

Using the testing dataset, the coefficient of determination (R^2^) was calculated for each curve, and the averaged coefficient of determination of the six curves for each input-output pair was obtained. The predictions were sorted according to their averaged coefficient of determination. The best, median, worst cases are plotted in Figure 8, and the corresponding actual and predicted material parameters are shown in Table 2. Perfect agreement is achieved for the best cases. For the median case, although the discrepancies in the constative parameters seem obvious, the six curves still have good matches. In the worst case, the results are still acceptable in terms of material parameters, and the actual and predicted stress-stretch curves follow the same trends.

**Table 2.**
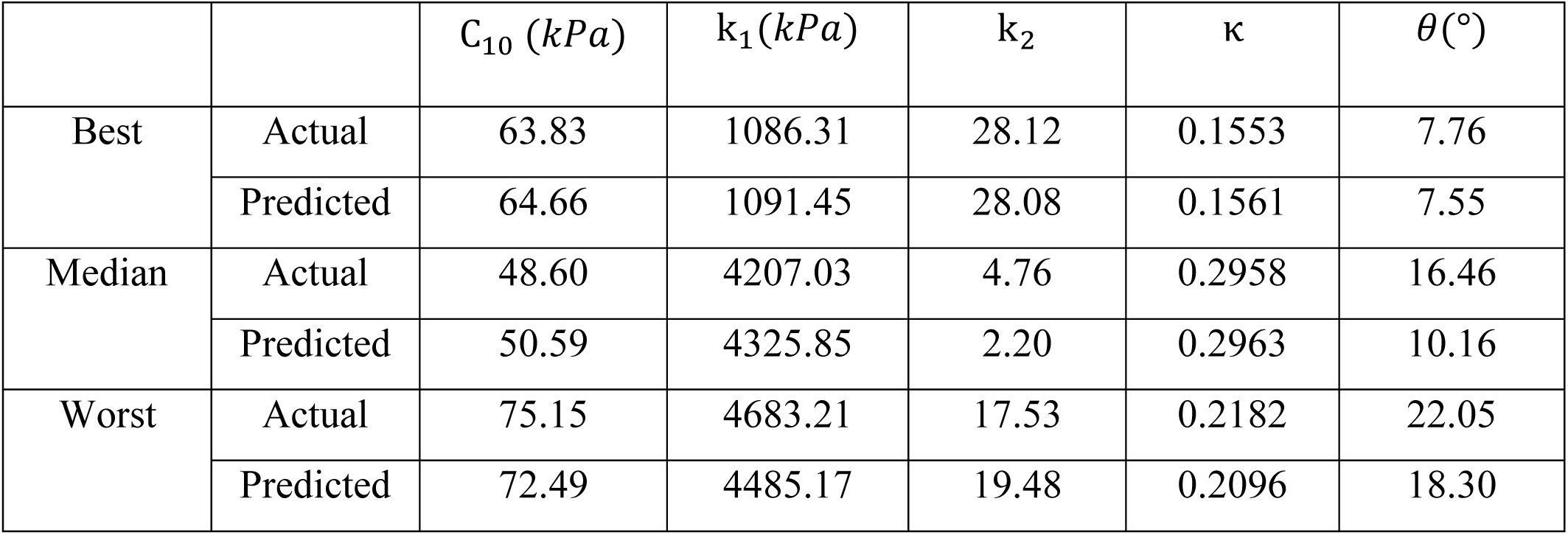
The actual and predicted material parameters for the best, median, worst cases.

## 4. DISCUSSION

Optimization-based inverse methods (Liu et al., 2017, 2018; Wittek et al., 2016; Wittek et al., 2013) have been extensively used for material parameter identification problems. Whether being dependent on FE simulations or not, these methods are computationally-expensive. Iterative computations limit the efficiency of these approaches, obstructing their clinical application. The proposed machine learning approach can fundamentally resolve the challenge on computation cost. The ML-model builds a direct linkage between the geometries and the material parameters, bypassing the iterative procedures. Once the ML-model is trained, it can be used to make predictions instantaneously and repeatedly, such that *in vivo* material parameter estimation on a high volume of patients in real-time becomes feasible. Although FE simulations are used to generate training, validation and testing datasets, which takes approximately 10 days – note that a similar amount of time is required to find the optimal material parameters for a single patient using nonlinear optimization (Wittek et al., 2016; Wittek et al., 2013). In terms of the predictions, the ML-model does not lose accuracy compared with the optimization-based methods developed by our group. The proposed ML-model was evaluated using additional testing data, where small discrepancies (NMAE) were achieved between the actual and ML-predicted material parameters. The close resemblance between the actual and predicted stress-stretch curves further demonstrates the high accuracy of the ML-predicted constitutive response.

The applications of machine learning techniques on the complex inverse mechanics problems can be traced back to the 1990s (Yagawa and Okuda, 1996), when neural networks were introduced to traditional mechanics fields for constitutive modeling (Ghaboussi and Sidarta, 1998) and elastic-plastic fracture mechanics (Theocaris and Panagiotopoulos, 1993). The pioneering work by (Huber and Tsakmakis, 1999a, b), determined some constitutive parameters from the spherical indentation data using neural networks. Because this classical problem can be characterized by the load-depth trajectory, some manually selected features (e.g. depth at a given load level) were sufficient. However, to determine material parameters of the aortic wall from medical image data, the 3D geometrical information has to be fully exploited, which cannot be done by using a few intuitive features. In our ML-model, the PCA effectively encodes the input complex geometries into the shape codes. Next, a neural network (24 inputs - 256 hidden units - 256 hidden units - 5 output units) with softplus activation function was utilized to establish the nonlinear mapping between the shape codes and the material parameters. The comparison between the softplus units and other units is illustrated in Figure 9. The softplus units outperformed the conventional sigmoid and hyperbolic tangent (tanh) units, the ReLU (Glorot et al., 2011) and its variant SELU (Klambauer et al., 2017). The softplus units lead to the lowest loss in the testing set and thus are more appropriate for this application.

**Figure 9.**
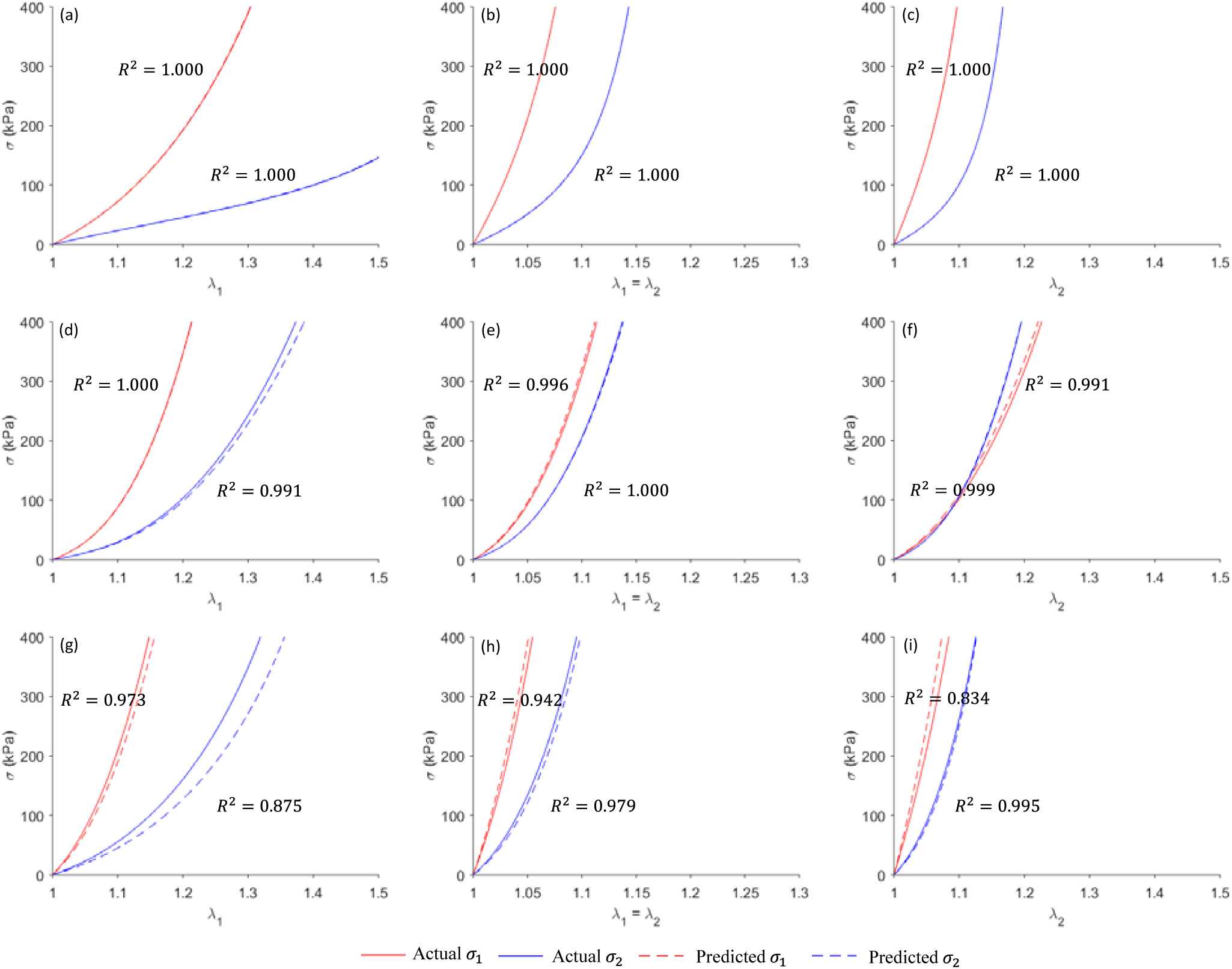
The actual and predicted stress-stretch curves for the best ((a), (b) and (c)), median ((d), (e) and (f)) and worst cases ((g), (h) and (i)).

**Figure 10.**
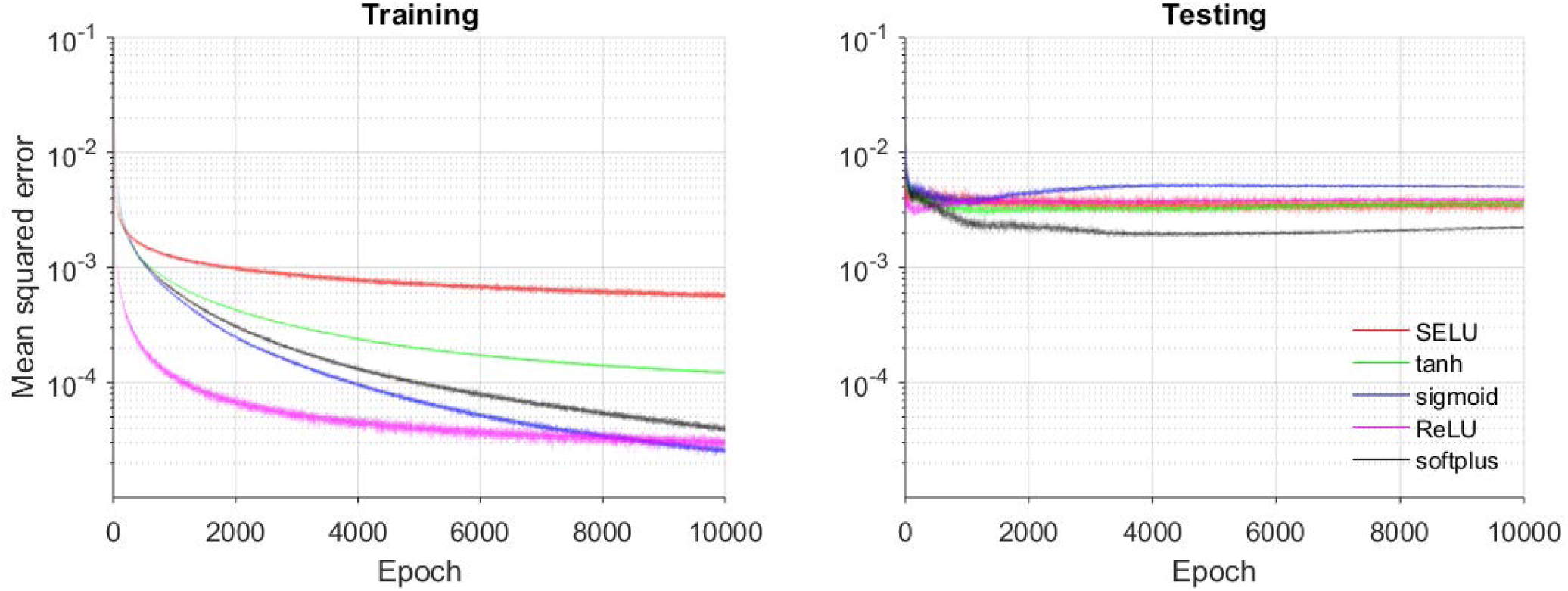
MSE loss function for training and testing using softplus and other units.

Since this paper only aims to demonstrate the feasibility of the proposed machine learning framework, the following assumptions and simplifications were used in data generation to expedite the FE simulations: (1) the branches of the aortic arch were trimmed off; (2) because of the partial volume effect, it is difficult to obtain the wall thickness from CT images, therefore, a uniform wall thickness at the systolic phase (1.5 mm) was assumed; (3) the systolic and diastolic pressure were fixed to 120mmHg and 80 mmHg, respectively. These limitations can be resolved in future work. The branches can be re-meshed using existing mesh processing method (Botsch et al., 2010), then encoded by additional shape codes. Using advanced magnetic resonance imaging (Dieleman et al., 2014), the wall thickness is measurable. The full 3D geometries at the two cardiac phases can be encoded using PCA, and therefore the thickness field is incorporated. To handle pressure variations, FE simulation data at a wide range of systolic and diastolic pressure levels can be generated, and the systolic and diastolic pressure can be simply included as two additional inputs to the neural network.

Although the feasibility of the ML-model is clearly shown, it is not ready for clinical application yet before more data become available. When a substantial amount of medical image data and experimental testing data are obtained, we can update the SSM space and the convex hull, from which a new large training dataset can be generated using the framework proposed in this study. The updated ML-model will be capable of accurately predicting the material parameters which may provide clinically relevant insights, i.e. serving as a basis for patient-specific rupture risk estimation (Martin et al., 2015). In case of a new patient with extreme aorta shape or material properties, which may cause unreliable prediction, a rejection option can be added in the ML-model as in (Bartlett and Wegkamp, 2008). The enhanced ML-model will avoid making predictions on uncommon cases.

## 5. CONCLUSION

We have proposed a novel ML approach to estimate the constitutive parameters of the aortic wall from *in vivo* loaded geometries at two cardiac phases with known blood pressures. The ML-model is comprised of an unsupervised shape encoding module and a supervised nonlinear mapping module. FE simulations were used to generate datasets for training, adjusting and testing the ML-model. This novel ML approach can expedite the procedure of *in vivo* material parameter identification: once the ML-model is trained, the material parameters can be estimated within one second.

## ACKNOWLEDGEMENTS

Research for this project is funded in part by NIH grant R01 HL104080 and HL127570. Liang Liang is supported by an American Heart Association Post-doctoral fellowship 16POST30210003.

